# On the origin of F-wave: involvement of central synaptic mechanisms

**DOI:** 10.1101/2023.06.12.544675

**Authors:** M. Görkem Özyurt, Filipe Nascimento, Robert M. Brownstone, Marco Beato

## Abstract

Neurophysiological methods are used widely to gain information about motoneuron excitability and axon conduction in neurodegenerative diseases. The F-wave is a common biomarker used to test motoneuron properties in the diagnosis of neurological diseases. Although the origin of the F-wave is a subject of debate, the most widely accepted mechanism posits that the F-wave is generated by the backfiring of motoneurons stimulated antidromically from the periphery. In this study, we developed an *ex vivo* mouse sciatic nerve-attached spinal cord preparation with sensory axons severed in which stimulation of the whole sciatic nerve or its tibial branch evoked responses with the electrophysiological signatures of F-waves. Manipulations of synaptic transmission by either removal of extracellular calcium or block of post-synaptic glutamate receptors abolished these responses. These results suggest that F-waves are mediated by spinal microcircuits activated by recurrent motor axon collaterals via glutamatergic synapses.

## Introduction

Neurological disorders that affect motor systems, such as Amyotrophic Lateral Sclerosis (ALS) and peripheral neuropathies, lead to substantial alterations in the properties of spinal motoneurons. In clinical neurophysiology, electromyographic (EMG) recordings are commonly employed to diagnose and understand the progression of such conditions. Electrophysiological parameters such as F-wave amplitude and latency, first described by Magladery and McDougal^1^, are important measurements of motoneuron and motor nerve excitability. Typically, F-waves have low and variable amplitude (2-5% of the maximal direct motor response), variable latency, and is subject to failures^2^. F-waves are often used to estimate motor axon conduction velocity and provide valuable information about a wide range of motor disorders, such as polyneuropathies and demyelinating conditions^2-4^.

The amplitude and latency of F-waves are both clinically useful properties. For example, a sensitive marker of abnormalities in lumbosacral radiculopathies^5,6^ is chronodispersion: the difference between minimal and maximal F-wave latencies in response to a given stimulus. And the amplitude, duration, and probability of occurrence of F-waves are used to gauge the excitability state of motor pools in a variety of central nervous system disorders^7-11^. For example, in the early stages of ALS, alterations in the excitability of motoneurons^12,13^ correlate with increased amplitude of the F-wave^11^.

Although F-waves are routinely used for diagnosing and understanding a variety of neurological disorders, the mechanisms underlying this delayed evoked response are not fully understood. The prevalent explanation for the origin of the F-wave is that the antidromic spikes elicited by electrical stimulation of motor axons lead to somatic depolarization in a variable minority of motoneurons, which in turn leads to the generation of a second spike that is propagated orthodromically and thus recorded as a low amplitude motor response (i.e. F-wave)^2,14^. While this is the generally accepted mechanism for the generation of F-waves, it is difficult to reconcile the concept of rebound spikes with the inactivation kinetics of sodium channels that would presumably prevent the generation (or propagation) of a second spike following the initial antidromic invasion. It is also notable that antidromic stimulation has been used to identify motoneurons in a variety of experimental systems *in vivo* and *in vitro*^15,16^, but the occurrence of a second “rebound” spike has never been reported.

In this report, we investigated an alternative explanation for the generation of F-waves. Studies using *in vitro* spinal cord preparations from mice have shown that motoneurons are synaptically connected with other motoneurons. Thus, antidromic activation of axons leads to excitation of other motoneurons via recurrent motor axon collaterals, a phenomenon known as recurrent excitation^16,17^. This excitation, mediated by glutamatergic transmission, can lead to orthodromic action potentials^16^. We thus hypothesize that recurrent excitation would be detected as a low-amplitude motor response in the muscle and could thus underlie F-waves. Since experiments in humans are not amenable to pharmacological manipulations, we tested this hypothesis using *ex vivo* preparations of neonatal mouse spinal cords with sciatic nerves attached. This preparation allowed us to evoke and record the equivalent of F-waves directly from the nerves while retaining the ability to use pharmacological manipulations. We present evidence that supports the hypothesis that F-waves result from recurrent excitation of motoneurons.

## Materials and Methods

### Animals

All the procedures were conducted in accordance with the Animal (Scientific Procedures) Act (Home Office, UK, 1986) and were approved by the UCL Ethical Committee, under project licence number PP2688499. *Ex vivo* experiments with nerve attached were performed on tissue obtained from female (n=7) or male (n=4) mice on postnatal days 1-3 (P1-P3). Patch clamp experiments on dorsal horn ablated spinal cords were performed on preparations female (n=4) or male (n=6) wild-type mice bred on a C57BL/6J background at P5-13.

### Dorsal horn ablated spinal cord preparations

We performed whole-cell recordings from labelled motoneurons in dorsal horn ablated preparations using the method we have previously reported ^18^. Briefly, P2-6 days old mice were anesthetized using isoflurane and we injected ankle dorsiflexor tibialis anterior (TA) or plantar flexor gastrocnemius (GS) muscles with Cholera Toxin Subunit B (CTB) conjugated Alexa Fluor 488 or 555 (1 μL and 0.2% wt/vol), respectively, to retrogradely label motoneurons. After 2-5 days, mice were anesthetised through intraperitoneal injection of a ketamine/xylazine mixture (at doses of 100 mg/kg and 10 mg/kg, respectively), and decapitated. The spinal cord was then dissected in ice-cold artificial cerebrospinal fluid (aCSF) containing (in mM) 113 NaCl, 3 KCl, 25 NaHCO_3_, 1 NaH_2_PO_4_, 2 CaCl_2_, 2 MgCl_2_, and 11 D-glucose and continuously bubbled with 95% O2 and 5% CO_2_. L3-L5 ventral roots-attached spinal cord was placed in agar block ventral side facing up, longitudinally cut using vibratome (HM 650 V, Microm, Thermo Fisher Scientific, UK) and incubated in a chamber with aCSF at 37°C for 30 mins, and kept at room temperature (∼21ºC) during electrophysiological recordings.

### *Ex vivo* spinal cord preparations with nerves attached

Animals were euthanized by cervical dislocation followed by decapitation. Mice were skinned and eviscerated with hindlimbs attached, keeping the spinal column intact. Following mid-thoracic spinal transection and vertebrectomy, the sciatic nerve, and triceps surae muscles were dissected in ice-cold aCSF. The spinal cord was gently lifted using a glass pipette to gain access to the dorsal side to perform full bilateral dorsal rhizotomies. Ventral roots were also cut, sparing L4 and L5. The common peroneal nerve and the tibial nerve branches were identified and dissected until the hip joint. The *ex vivo* spinal cord with sciatic nerve attached (**Figs 2A and 3A**) was then immersed in aCSF which was continuously bubbled and kept at room temperature (∼21ºC) before being used for electrophysiological recordings.

### Electrophysiology

Patch clamp recordings were performed using an Axopatch 200B amplifier (Molecular Devices, Sunnyvale), low pass filtered at 5 kHz, digitised at 50 kHz with a Digidata 1440A A/D board (Molecular Devices), and acquired with Clampex 10 software (Molecular Devices). *Ex vivo* nerve recordings were performed using an NPI Ext-02F amplifier (NPI electronic GmbH) and signals were bandpass filtered at 1Hz – 10 kHz. Borosilicate glass (GC150F, Harvard Apparatus) electrodes for recording and stimulation were pulled with a Flaming-Brown puller (P1000, Sutter Instruments), cut to ∼1.5x the root or nerve diameter for stimulation and nerve recordings, attached to a head stage (Narishige), filled with regular aCSF, then attached to roots by suction. Electrical current was delivered using a DS3 isolated current stimulator (Digitimer). Patch recordings were performed using glass pipettes (∼1–4 MΩ resistance) filled with intracellular solution containing (in mM): 125 K-gluconate, 6 KCl, 10 HEPES, 0.1 EGTA, 2 Mg-ATP, pH 7.3 with KOH, and osmolarity of 290–310 mosmol/kgH2O. Membrane potentials were reported without correcting the junction potential for the intracellular solution (∼15 mV).

*In vitro* spinal cords were placed in a chamber continuously perfused with aCSF at room temperature and imaged using Eclipse E600FN Nikon microscope (Nikon, Japan). Infrared differential interference contrast images were captured with a digital camera (Nikon, DS-Qi1Mc), while fluorescence was detected through a laser scanning confocal unit (D-Eclipse C1, Nikon) equipped with two diode laser lines (λ = 488 and 561 nm). The stimulus intensity was defined by the threshold intensity, which is the minimum intensity that generated excitatory postsynaptic potentials (EPSP) reliably in motoneurons. Both L4 and L5 ventral roots were stimulated with 2-3x threshold intensity. Labelled motoneurons (n=11 TA, n=8 LG) were patched (**Fig 1A-B**) and stimulus evoked responses were recorded in the current-clamp configuration. The occurrence of antidromic and orthodromic spikes at different resting potentials were tested in 25 unlabelled motoneurons (**Fig 1C**).

**Figure 1.**
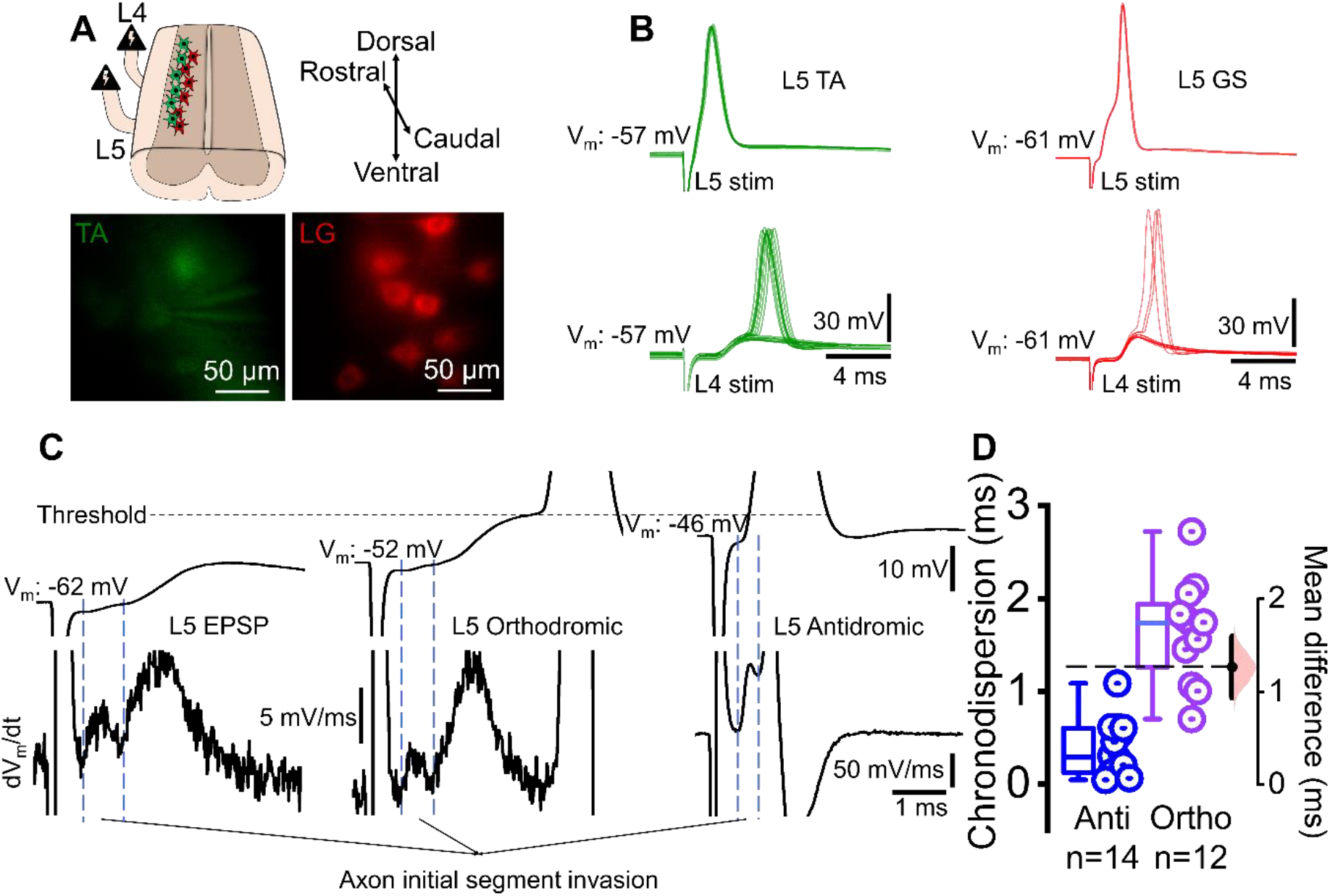
Ortho-and antidromic action potentials generated in motoneurons following stimulation of ventral roots. **A)** Drawing of the dorsal horn ablated spinal cord preparation with L4 and L5 ventral root stimulation (upper) and representative images of labelled tibialis anterior (TA) and gastrocnemius (GS) motoneurons. **B)** Current clamp recordings of ventral root evoked responses in one TA (left, 5 sweeps) and one GS (right, 2 sweeps) motoneuron, showing homosegmentally-evoked antidromic spikes (upper). At resting membrane potential, orthodromic spikes and occasional spike failures with visible EPSPs were recorded in response to adjacent root stimulation (lower, 14 sweeps for TA and 9 sweeps for GS). **C)** Current clamp recordings of L5 ventral root evoked responses in unlabelled dorsolateral motoneurons (upper panels) showing subthreshold excitatory postsynaptic potential (EPSP), near threshold EPSP-evoked orthodromic spike, and antidromic spike. Lower panels show derivatives showing antidromic axon initial segment spikes even in the absence of somatic antidromic spikes. **D)** Plot on the right shows chronodispersion of antidromic and orthodromic spikes. Each point indicates a motoneuron and the graph on the right is the estimation plot.

The spinal cord–sciatic nerve *ex vivo* preparation was placed in a chamber with aCSF continuously perfused at 5 mL/min. Either the tibial or whole sciatic nerve was stimulated, preferentially from a distal location along the isolated nerve, with the nerve recordings being performed with a suction electrode positioned on a proximal section of the sciatic nerve (see **Figs 2A** and **3A**). The nerve was stimulated every 15 seconds with 0.2-0.5 ms pulse width. Stimulus intensity was set at around 10% above the intensity that generated the maximum early direct response measured from the sciatic nerve. We refer to this early response as the “direct response,” recognising that it is not equivalent to the direct response recorded clinically from the muscle as sensory axons will also be stimulated; we used this direct response as a reference to test the stimulation efficiency throughout the experiment. That is, we constantly monitored this direct response and the F-wave from the sciatic nerve (or tibial branch), and the experiments were terminated if the reduction in the direct response exceeded 20%. Throughout the text, we have reported the latency and chronodispersion (the difference between the maximum and minimum F-wave latency) of the F-wave response as the mean ± standard deviation usually obtained from ∼10 or more individual traces. The conduction velocity (CV) was calculated as follows; the distance from the stimulation site to the cord and back to the recording (in mm) site was divided by the latency (ms) of the F-wave. Manipulation of the synaptic excitability was done by lowering extracellular Ca^2+^ to 0 mM or exogenous application of the α-amino-3-hydroxy-5-methyl-4-isoxazolepropionic acid (AMPA) receptor antagonist - 3-Dioxo-6-nitro-1,2,3,4-tetrahydrobenzo[f]quinoxaline-7-sulfonamide disodium salt (NBQX) - at a concentration of 6 μM that is effective to block the AMPA receptor mediated currents^19^. Graphs showing the mean size of the F-wave and chronodispersion were plotted using OriginPro 2021 (OriginLab Corporation, Northampton, MA), and estimation plots^20^ were plotted with MATLAB R2022a (Mathworks, Natick, MA).

## Results

We recorded motoneurons innervating either the LG or TA muscle in dorsal horn ablated spinal cords and stimulated ventral roots in the same or adjacent segment to the recorded motoneuron (**Fig 1A**). In all cases, an antidromic spike was observed following same segment stimulation (**Fig 1B - upper traces**), but the subsequent EPSP was never sufficient to evoke an orthodromic spike following the antidromic one. On the contrary, stimulation of the adjacent segment invariably resulted in an EPSP that could exceed threshold and give rise to an orthodromic spike (**Fig 1B - lower traces**).

We next tested whether we could prevent antidromic invasion of the soma by hyperpolarizing the motoneuron, and if in so doing, an orthodromic spike could occur. Indeed, in 14 out of 25 recorded motoneurons with antidromic spikes, hyperpolarization (between -60 and -75 mV) by direct current injection prevented somatodendritic (SD) antidromic spike but not the axon initial segment (IS) spike (**Fig 1C**, left), visible as a peak in the first derivative of the voltage trace (bottom row of **Fig 1C**, left). This IS spike was followed by an EPSP that did not reach threshold. Reducing the injected current (**Fig 1C**, middle) enabled the EPSP to reach threshold and generate an orthodromic spike. But when the motoneuron was held at its resting membrane potential (**Fig 1C**, right) the occurrence of the antidromic spike prevented the occurrence of the orthodromic one. In 22 of these motoneurons, we also compared the chronodispersion of the anti and orthodromic spikes recorded and found a greater degree of chronodispersion for the orthodromic spikes than for the antidromic ones (**Fig 1D**, 1.66 ± 0.55 ms vs. 0.39 ± 0.29 ms). This was not unexpected, given the EPSP reaches the threshold at different times during its rising phase.

We reasoned that in an intact system these orthodromic spikes could give rise to a delayed motor response propagating along the nerves and thus account for F-waves. We next tested if we could evoke and measure a motor response that would have the electrophysiological signatures of the F-wave (**Fig 2A**). We stimulated the whole sciatic nerve (P1-3, n=5) and detected a large, early direct response at a more proximal sciatic site (**Fig 2B**). Following this direct response, in all the animals tested, we detected a subsequent response with a latency of 29.1 ± 5.6 ms, with 3.9 ± 0.8 ms chronodispersion (CV: 0.58±0.08 m/s, in the n=3 cords in which the orientation of the nerve allowed accurate length determination). Our CV estimate is similar to the reported neonatal mice motor axon CV^21^. This second response could be regarded as an *ex vivo* analogue of the F-wave that is usually measured through EMG since it: 1) followed an initial direct motor volley, 2) did not arise from sensory axon mediated reflexes, and 3) was variable in size, shape and latency (see individual sweeps in **Fig 2B-E**, chronodispersion in **Fig 2G**). Since these features matched the characteristics of the F-wave studied in clinical neurophysiology, we conclude that in our *ex vivo* neonatal mouse preparations, we were able to successfully evoke and measure F-waves from nerves.

Given our postulate that orthodromic action potentials lead to F-waves (**Fig 1**) and given that we could record an F-wave equivalent (**Fig 2**), we were now able to ask whether F-waves are generated by synaptic mechanisms. That is, could we abolish the F-wave response by manipulating synaptic transmission at either pre- or post-synaptic sites?

We first lowered synaptic release probability by removing Ca^2+^ from the extracellular solution. Doing so did not reduce the direct response (∼4% increase compared to control condition), but completely abolished the F-wave (**Fig 2C**). This effect was reversible: following reequilibration with 2 mM extracellular Ca^2+^, the F-wave recovered (**Fig 2D**), indicating that blocking pre-synaptic transmitter release by removing Ca^2+^ is sufficient to prevent the generation of the F-wave.

Since Ca^2+^ removal could also modulate motoneuron excitability^22^, we next blocked post-synaptic receptors. Since glutamate (AMPA) receptors are known to mediate recurrent excitation between motoneurons^16^ and blocking them would not affect motoneuron excitability^16^, we used a selective AMPA antagonist, NBQX^19^. We found that exogenous application of NBQX (6 μM) completely suppressed F-waves without affecting the direct motor response (∼2% increase compared to control condition) (**Fig 2E**). That is, we found that F-waves can be completely suppressed by either impairing the synaptic release machinery (reducing the probability of release by removing Ca^2+^) or by blocking post-synaptic receptors (**Fig 2F**). These results can be explained if F-waves are generated by recurrent synaptic connections, but not if they are generated by rebound spikes.

In clinical studies the F-wave is usually evoked by stimulating a single branch of the sciatic nerve. We therefore repeated our experiments stimulating only the tibial nerve and recording from the sciatic nerve using the same stimulation paradigm as above (**Fig 3A**). Similar to the previous set of experiments, following tibial nerve stimulation, we detected an initial direct response, followed by a long latency event (37.1 ± 6.4 ms, CV: 0.42±0.01 m/s for n=2), smaller in size and with clear chronodispersion of 5.3 ± 1.3 ms (P1-3, n=6, **Fig 3B and F**). This shows that stimulation of a single branch of the sciatic nerve, in a configuration similar to that used during clinical tests, is sufficient to evoke the *ex vivo* analogue of the F-wave.

**Figure 2.**
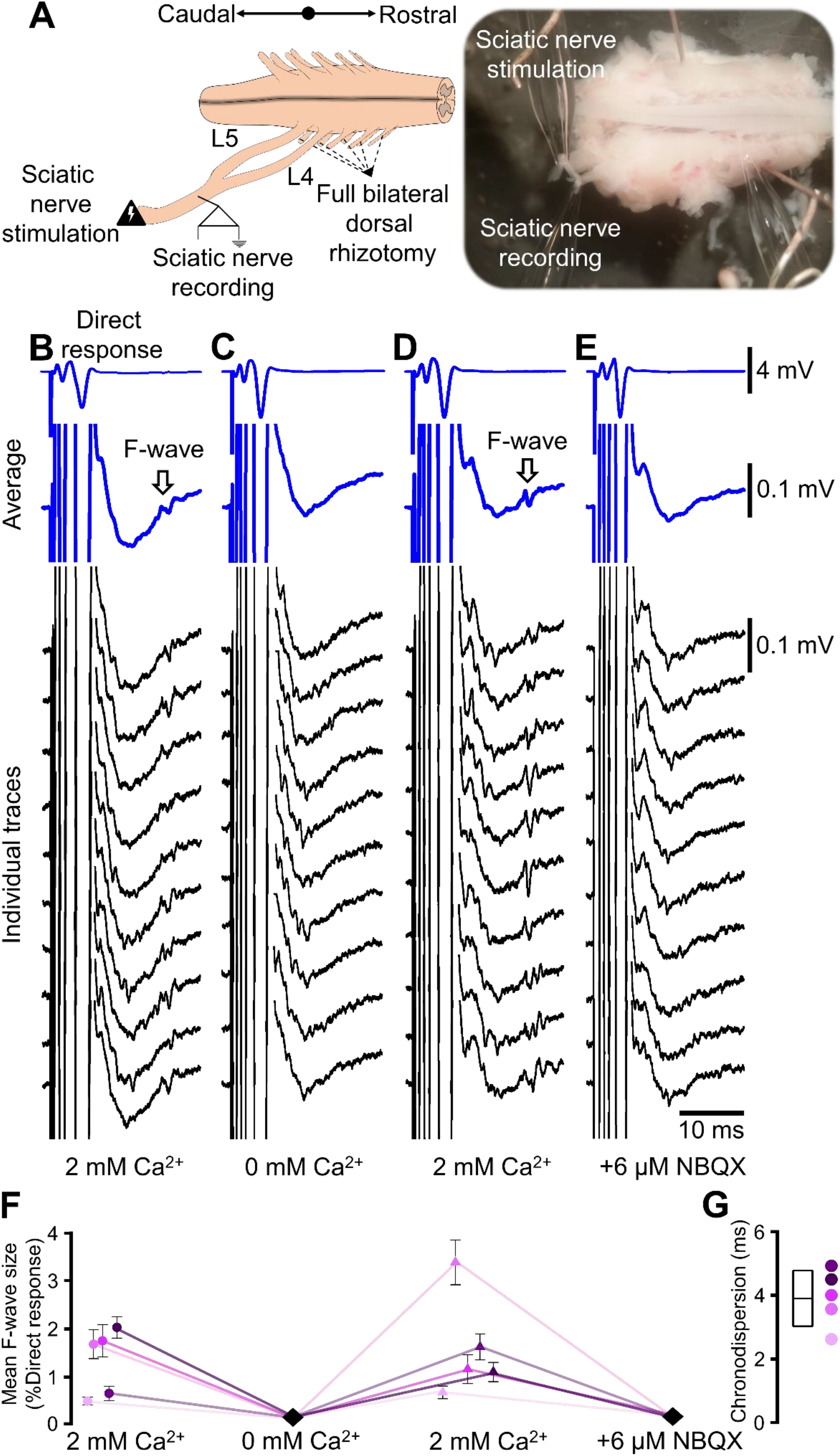
F-waves recorded from neonatal mouse spinal cord – sciatic nerve *ex vivo* preparations following sciatic nerve stimulation depend on central synaptic release mechanisms. **A)** Schematic (left) and representative photograph (right) of the isolated spinal cord – sciatic nerve preparation, depicting the sites of stimulation (distal sciatic nerve) and recording (proximal sciatic nerve). **B)** Example of averaged (blue) and individual (black) traces obtained from a 2-day-old male mouse pup. Direct responses are amplified in the second averaged trace to visualise the long latency F-wave, indicated by arrows. **C)** Ca^2+^-free aCSF abolished the F-wave, that **D)** was restored upon reperfusion with a solution containing 2 mM Ca^+2^. **E)** Perfusion with the AMPA receptor antagonist NBQX, also abolished the F-wave. **F)** Plots show mean F-wave amplitudes in each animal (different colours) in different conditions and **G)** chronodispersion of F-wave in control 2 mM Ca^2+^ condition (mean ± standard deviation).

**Figure 3.**
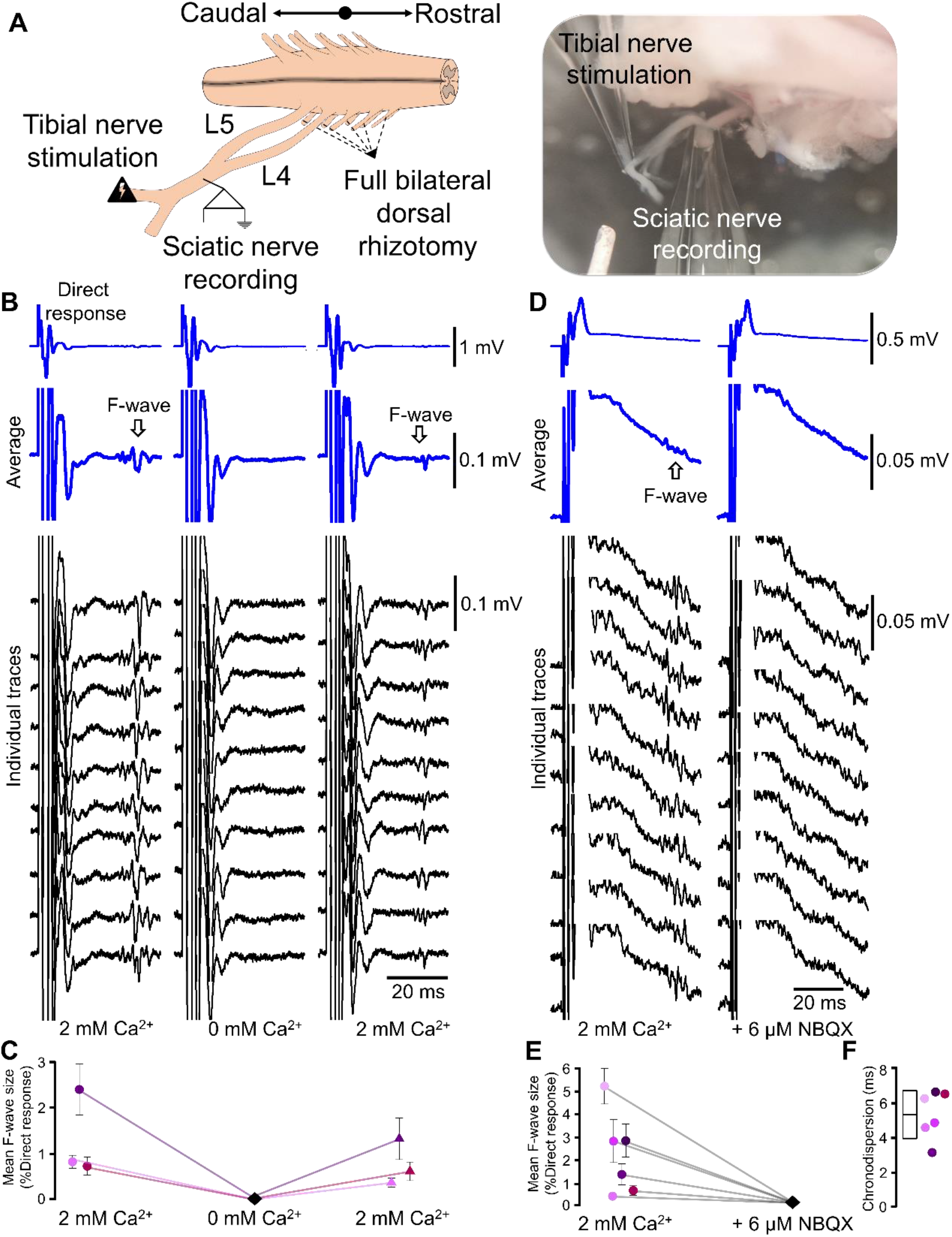
Stimulation of the tibial nerve also elicits F-waves that depend on central glutamate release. **A)** Drawing (left) and photograph (right) of the isolated spinal cord – sciatic/tibial nerve neonatal mouse preparation, depicting stimulation (tibial nerve) and recording (sciatic nerve) sites. **B and C)** Representative averaged (blue) and individual (black) traces obtained from a 2-day-old mouse pup, showing the direct motor responses (top row). These traces are increased in amplification (2^nd^ row) to show the F-wave responses (indicated by arrows). The F-wave recorded in the presence of 2 mM extracellular Ca^2+^ (left) was abolished in Ca^2+^-free aCSF (middle) and recovered following reperfusion of 2 mM Ca^2+^ solution (right). **D and E)** The AMPA receptor antagonist, NBQX (6 μM), abolished the F-wave (different preparations from B). **F)** F-wave chronodispersion at control Ca^2+^ levels (mean ± standard deviation).

In order to confirm the synaptic identity of the F-wave recorded at the sciatic nerve following tibial nerve stimulation, we repeated the experiments blocking either pre-synaptic release or post-synaptic receptors. Blocking pre-synaptic release by removing Ca^2+^ did not affect the direct response (∼1% increase compared to control condition) but resulted in complete suppression of the F-wave, which was then restored upon reapplication of normal aCSF containing 2 mM of Ca^2+^ (**Fig 3B, C**). Similarly, blocking AMPA receptors with NBQX (6 μM) completely abolished the F-wave without affecting the initial motor response (∼4% reduction compared to control condition, **Fig 3D, E**). Altogether, these experiments showed that the clinically-relevant analogue, stimulation of a branch of the sciatic nerve (i.e. tibial nerve), is sufficient to generate the F-wave *ex vivo*, and that the F-wave is abolished by blocking either transmitter release or post-synaptic AMPA receptors. That is, the F-wave results from synaptic activity.

## Discussion

We used an *ex vivo* neonatal mouse spinal cord preparation with the sciatic nerve intact and showed that antidromic activation of motor axons from either the whole sciatic nerve or its posterior branch (tibial nerve), can evoke relatively long latency responses in the sciatic nerve. This response has the fundamental electrophysiological signatures of the F-wave^2^. We demonstrated that this F-wave is abolished by blocking synaptic transmission by either removing extracellular Ca^2+^ or by blocking post-synaptic glutamate receptors. These observations indicate that the F-wave is synaptically generated by glutamatergic spinal microcircuits activated by synchronous motoneuron firing.

In clinical settings, low intensity stimulation of a mixed nerve generally evokes an initial H-reflex in the muscles. The gradual increase in the intensity initially results in a larger H-reflex and in activation of motor axons such that a direct motor response (M-response) is generated. However, further increment in intensity leads to an increase in the M-response while the H-reflex decreases in amplitude until it is completely abolished, due to collision of the antidromic and orthodromic motor volley^23^. After this stage, an F-wave appears, characterised by low amplitude, variable shape, and high jitter. F-waves result from activation of motor axons, possibly those innervating fast twitch muscle fibres^24,25^. Sensory afferent activation is not required, as the F-wave can still be obtained in deafferented patients or in animals with severed dorsal roots^26,27^. However, despite being widely used, the physiology of the F-wave is not yet understood.

The commonly accepted idea behind the F-wave is that stimulation of motor axons leads to re-excitation of the somatodendritic membrane that subsequently results in the stimulated motoneuron re-firing, giving rise to a “rebound” F-wave^14,28-30^ (**Fig 4**, left side). The variable shape of the F-waves across trials was attributed to different motoneurons producing the rebound firing in different trials^31^.

**Figure 4.**
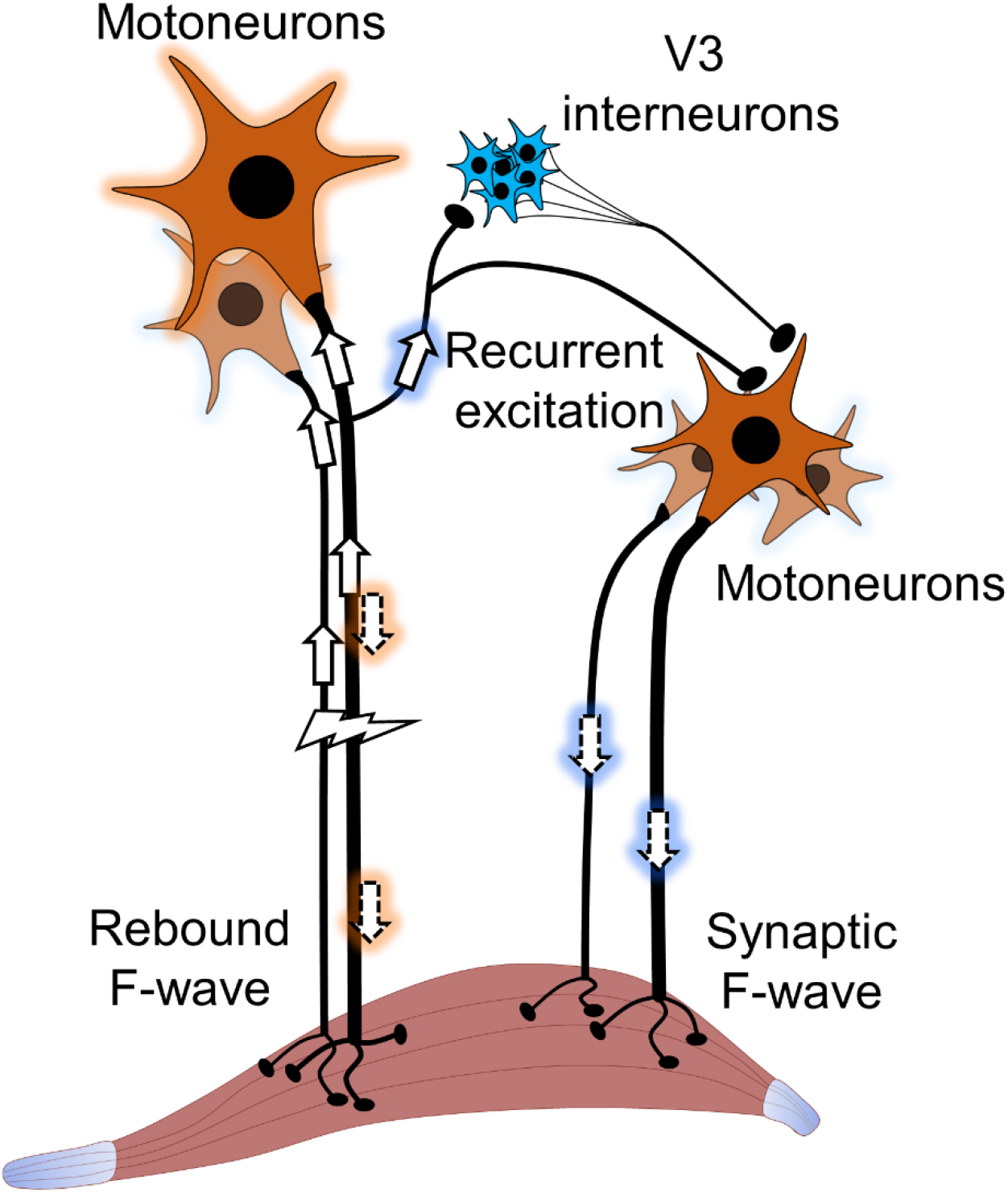
Possible mechanisms behind the F-wave. Classical explanation is that the stimulation of motor axons antidromically leads to the re-activation of some of the motoneurons, which then triggers a “rebound” response (left). This response is recorded in the muscle as the F-wave. The mechanism proposed here is based on central synaptic glutamate release. The stimulation of motor axons leads to the generation of action potentials in some motoneurons through recurrent excitatory collaterals, activating either mono-or disynaptic excitatory microcircuits (V3 interneurons or other unknown pathways), resulting in a “synaptic” F-wave. The motoneurons generating the synaptic F-wave are also stimulated but for representative purposes this is not shown on the figure.

Here we show that, at least in *ex vivo* preparations, F-waves result from efferent-triggered central glutamate release. Antidromic invasion of motor axon collaterals can activate various circuits, including inhibitory Renshaw cell recurrent loops^32^ as well as multiple excitatory microcircuits. One of these excitatory loops is made by motor axon collaterals forming glutamatergic synaptic connections with other ipsilateral motoneurons^16,18,33,34^. Recurrent motor axon collaterals also activate a ventrolateral population of V3 interneurons, which in turn form glutamatergic synapses with motoneurons^17^. Of note, motoneuron to motoneuron connections are maintained beyond the neonatal stage and span beyond a single spinal segment^16^. These connections are ten times greater in large post-synaptic motoneurons innervating fast twitch muscle fibres compared to small ones that innervate slow twitch fibres ^16,18^. As such, so synaptically-generated F-waves, as shown here, would be expected to be predominant in large motoneurons – a suggestion compatible with the clinical observation that F-waves are primarily generated by larger motor units^24,25^.

In our in vitro conditions (**Fig 1**), as well as in all the in vivo recordings we are aware of, somatic invasion of the antidromic spike prevents the generation of an early orthodromic spike. In fact, an SD spike will prevent a second axonal spike generated by intracellular current injection unless the motoneuron is hyperpolarised to a point where there is a significant IS-SD delay^35^. If the motoneuron is relatively hyperpolarized at the time of arrival of an antidromic impulse (evoked by peripheral stimulation), the IS spike may not activate the SD membrane^36^. In this case, the recurrent EPSP could be sufficient to generate an orthodromic spike that, if sufficiently delayed beyond the absolute refractory period of the axon^35^, would then propagate towards the muscle (**Fig 4**, right side). With F-waves being produced via orthodromic (synaptic) activation, they will be particularly sensitive to membrane voltage because the voltage would need to be hyperpolarised sufficiently to block the SD spike, and yet not so much that the EPSP does not reach threshold. In fact, it has been suggested that F-waves reflect the state of motoneuron inhibition^8^, a condition that could lead to SD spike failure. In awake animals, motoneuron membrane potential fluctuates, meaning that at the given moment of stimulation, a variable pool of motoneurons will participate in the F-wave. That is, fluctuations of membrane voltage could account for both the observed variability in amplitude and the chronodispersion because the stochastic nature of these events would make it unlikely that the same motoneurons (with the same conduction velocities) would be recruited from trial to trial.

Could understanding the physiology of F-waves provide any further insight into the pathophysiology of disease? For example, in people with ALS, the F-wave has reduced persistence but increased amplitude, latency and chronodispersion. Each of these effects can be explained by “rebound” action potentials, as well as by orthodromic synaptic activation. But if synaptic activation is necessary for F-waves, then perhaps the recurrent excitation of slow motoneurons as well as inhibitory inputs (to hyperpolarise motoneurons) both increase (supplementing a homeostatic response^37^) as the disease progresses. That is, the F-wave may be sensitive to changes in synaptic inputs to motoneurons across the course of the disease.

In summary, our results indicate that the F-wave is synaptically-mediated by recurrent excitation through motor axon collaterals. Our findings provide further evidence that the F-wave reflects not only the excitability of motoneurons but also their synaptic connectivity patterns. Both of these parameters may be affected in neurological disorders and may thus impact the F-waves.

## Acknowledgements

We would like to thank the members of Beato and Brownstone labs and acknowledge the contribution of funding agencies to support this study. MGO: Royal Society Newton International Fellowship NIF\R1\192316, FN: Sir Henry Wellcome Postdoctoral Fellowship 221610/Z/20/Z, RMB: Medical Research Council Research Grant MR/V003607/1, MB: Biotechnology and Biological Sciences Research Council Research Grant BB/S005943/1. RMB is a co-founder and is on the board of Sania Therapeutics, Inc and consults for Sania Rx Ltd.

